# Biological S^0^ reduction at neutral (pH 6.9) and acidic (pH 3.8) conditions: Performance and microbial community shifts in a H_2_/CO_2_-fed bioreactor

**DOI:** 10.1101/2024.05.11.592705

**Authors:** Adrian Hidalgo-Ulloa, Charlotte M. van der Graaf, Irene Sánchez-Andrea, Jan Weijma, Cees J.N. Buisman

## Abstract

Sulfidogenesis is a promising technology for the selective recovery of chalcophile bulk metals (e.g. Cu, Zn, and Co) from metal-contaminated waters such as acid mine drainage (AMD) and metallurgy waste streams. The use of elemental sulfur (S^0^) instead of sulfate (SO_4_^2-^) as electron acceptor reduces electron donor requirements four-fold, lowering process costs, and expands the range of operating conditions to more acidic pH. We previously reported autotrophic S^0^ reduction using an industrial mesophilic granular sludge as inoculum under thermoacidophilic conditions. Here, we examined the effect of pH on the S^0^ reduction performance of the same inoculum, in a continuously fed gas-lift reactor run at 30 °C under neutral (pH 6.9) and acidic (pH 3.8) conditions. Steady-state volumetric sulfide production rates (VSPR) dropped 2.3-fold upon transition to acidic pH, from 1.79 ± 0.18 g·L^-1^·d^-1^ S^2-^·to 0.71 ± 0.07 g·L^-1^·d^-1^ S^2-^· Microbial community analysis via 16S rRNA gene amplicon sequencing showed that at pH 6.9, the S^0^-reducing genera *Sulfurospirillum, Sulfurovum, Desulfurella*, and *Desulfovibrio* were present at the highest relative abundance, while at pH 3.9 *Desulfurella* dominated the sequenced reads. The detection of acetic acid and the relative abundance of *Acetobacterium* at pH 6.9 pointed towards acetogenesis, explaining the dominance of the heterotrophic genus *Sulfurospirillum* in this H_2_ and CO_2_–fed bioreactor.

## 1. Introduction

Metal removal from metalliferous waters such as acid mine drainage and hydrometallurgical streams through metal sulfide precipitation is advantageous over more commonly used chemical neutralization methods, as it enables pH-dependent selective metal recovery at sufficient purity for recycling (Lewis, 2010). Microbial sulfide production (biosulfidogenesis) is a preferred source of hydrogen sulfide (H_2_S), as it can be carried out on-site and modified to meet process demands (Johnson and Sánchez-Andrea, 2019). Although biosulfidogenic processes have been commissioned on an industrial scale (Adams et al., 2008; Huisman et al., 2006), predominantly based on sulfate (SO_4_^2-^) as the electron acceptor, the technology is not widely used in the hydrometallurgical industry, partly due to the operational expenditure (OpEx) related to substrate requirements (Sun et al., 2020a). Substrate utilization can be lowered by using elemental sulfur (S^0^) instead of SO_4_^2-^ as electron acceptor, as this enables a theoretical fourfold decrease in the electron donor consumption for generation of an equimolar amount of H_2_S (Florentino et al., 2016b).

Further process optimization and reduction of the OpEx and CapEx (capital expenditure) can be achieved by integrating biosulfidogenesis and metal recovery in one reactor unit, where H_2_S is produced in the hydrometallurgical process waters (Kumar et al., 2021). Given the frequent high acidity (pH < 4) and, in some cases, elevated temperatures (40 – 80 °C) of these waters, which arise from the upstream processing of the material, successful integration of sulfidogenesis and metal precipitation necessitates a microbial community adept at surviving in such extremophilic conditions.

We previously reported a S^0^-reducing continuous gas-lift bioreactor operated at thermoacidophilic conditions (pH 3.6, 60°C), using a neutrophilic industrial granular sludge as inoculum, and fed with H_2_ and CO_2_ as sole electron donor and carbon sources, respectively (Hidalgo-Ulloa et al. 2023). Under these conditions a maximum VSPR of 270 mg H_2_S·L^-1^·d^-1^ was achieved. This is up to fivefold lower than those obtained in other studies at mesophilic temperatures, both at acidic (pH 6.5 - 2.1) (Sun et al., 2020b) and neutral pH (Sun et al., 2018; Zhang et al., 2018b). Although differences in system configuration do not allow direct comparison, previous studies also reported higher VSPR under mesophilic compared to thermophilic conditions (Azabou et al., 2007; Segerer et al., 1985; Takai et al., 2003). Thermophilic conditions can furthermore lead to operational complications such as bioreactor corrosion and unintended formation of secondary minerals, causing the re-precipitation of valuable leached elements and the decrease of the metal recovery yield (Batty and Rorke, 2006; Hedrich et al., 2018). Therefore, focusing on process optimization at mesophilic temperatures presents an opportunity to improve the VSPR and enhance the overall process design.

Therefore, we followed up on our previous study by investigating reactor performance at mesophilic temperature (30 °C) at both, neutral (6.9 ± 0.1) and acidic pH (3.8 ± 0.1), using the same neutrophilic granular sludge industrial as inoculum. By comparing the VSPR at steady-state achieved in this study with those achieved under thermoacidophilic conditions, we aimed to identify possible limitations of our system. Furthermore, we investigated changes in the microbial community composition through 16S rRNA gene amplicon sequencing in the two pH regimes.

## 2. Materials and methods

### 2.1. Reactor configuration and inoculum preparation

A glass gas-lift reactor with a working volume of 4 L was inoculated with wet granular sludge from a SO_4_^2-^-reducing bioreactor with low methane production at the industrial chemical plant Getec park (Emmen, the Netherlands) (Hulshoff et al., 2001). The reactor was operated at 30 °C and supplied with low phosphate mineral media (Hidalgo-Ulloa et al., 2022), with H_2_ and CO_2_ as sole electron and carbon donors. Influent media was continuously sparged with N_2_ (O_2_ < 0.5 ppmv, Linde Gas Benelux B.V., the Netherlands). Settleable solids and suspended biomass were retained using a 1.1 L glass settler. Further description of the reactor configuration and equipment used is provided elsewhere (Hidalgo-Ulloa et al. 2023). The VSPR was estimated from the change in sulfide concentration 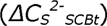 in the gas effluent scrubber (5M NaOH) over time (*Δt _(n+1,n)_)* (eq. 1), expressed in g·L^-1^·d^-1^ S^2-^.

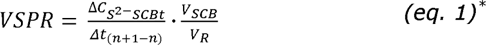

Upon start-up, mineral media (3 L) was added to the reactor. After sparging with N_2_ gas for 1 h (25 mL·s^-1^), N_2_ gas was substituted with a mixture of H_2_ (>99.999%, Linde Gas Benelux B.V) and CO_2_ (>99.99%, Linde Gas Benelux B.V) with a gas rate equal to the initial operating conditions (1 h). Concurrently, 10 g S^0^·L^-1^ of biological elemental sulfur (henceforth S^0^) was added to the reactor.

Prior to inoculation, 400 g (32 g dry weight) of the industrial wet granular sludge (henceforth Emmen sludge) was suspended in 500 mL of demineralized water, and the pH of the suspension was adjusted to 6.9 with 1 M H_2_SO_4_. The sludge suspension was sparged with N_2_ (25 mL·s^-1^) for 60 minutes before inoculation of the reactor.

After initial operation in batch mode for six days, operation was switched to continuous mode at a constant hydraulic retention time (HRT) of 2.5 days. Operational conditions were maintained, except during instances of technical disruptions (Supplementary information, S.I.1). During the continuous operation, S^0^ was added to the reactor in batch through a feed port. The amount of S^0^ supplied was based on a mass balance over the H_2_S produced. After 38 days of operation, we identified ammonium (NH_4_ ^+^) deficiency in the reactor; thus, we increased the concentration of (NH_4_)_2_SO_4_ from 0.55 mM to 2.53 mM to satisfy the microbial demand. The origin and preparation of the S^0^ used is described in detail by Hidalgo-Ulloa et al. (2023).

This research intended to examine the sulfidogenic capacity of the granular sludge at neutral (6.9 ± 0.1) and acidic (3.8 ± 0.1) pH. Operation started at neutral pH, during which a 0.1 M NaOH solution was used to maintain a constant pH. Once steady-state^†^ conditions were reached, the reactor pH was decreased to 3.8 using a 0.1 M H_2_SO_4_ solution. After the initial 14 days at pH 3.8, the pH control solution was replaced with 0.1 M HCl to limit the contribution of SO_4_^2-^ reduction to the VSPR. Additionally, different influent H and CO flow rates were tested. The reactor was initially fed with 2.8 L·h^-1^ of H_2_ and 0.7 L·h^-1^ of CO. The H inflow rate was increased to 5.6, 11.2, and 28 L·h^-1^, and then lowered to 8 L·h^-1^ until reaching steady-state. Likewise, the inflow rate CO_2_ was increased to 2 L·h^-1^ and afterward decreased to 0.7 L·h^-1^ until reaching steady-state.

### 2.2. Microbial community analysis

The microbial community was analyzed through 16S rRNA gene amplicon sequencing. Triplicate samples were collected from the anaerobic sludge and the washed S^0^, both of which were harvested by centrifugation and stored at -20 °C until further processing. The anaerobic sludge, collected from the Emmen plant in January 2018, was kept in a 10 L container at 4 °C. In March 2020, samples from this batch were prepared for DNA extraction. Samples for microbial community analysis were taken in triplicate during reactor operation on days 24, 59, 73, 101, 118 and 130. Reactor sampling, DNA extractions, PCR amplification, library preparation and sequencing were performed as described previously (Hidalgo Ulloa et al. 2023). The V4-V5 region from the 16S rRNA gene was amplified using PCR with barcoded revised Earth Microbiome Project (EMP) primers: 515F (GTGTGYCAGCMGCCGCGGTAA) and 806R (CCGGACTACNVGGGTWTCTAAT) (Thompson et al., 2017).

Paired-end amplicon sequences were processed using NG-Tax 2.0 on the Galaxy platform (https://ngtax.systemsbiology.nl) (Poncheewin et al., 2020) as described previously (Hidalgo Ulloa et al. 2023). Taxonomy was assigned using the SILVA SSU rRNA reference database v138 (Quast et al., 2013; Yilmaz et al., 2014), and sequences were further analyzed with R (Core Team, 2021) in RStudio, with the phyloseq (McMurdie and Holmes, 2013), microbiome (Lahti and Shetty, 2017), ggplot2 (Wickham, 2008), ggpubr (Kassambara, 2020) and dplyr packages in the tidyverse (Wickham et al., 2019). Alpha diversity was calculated on rarefied data (sample size 22050). Beta diversity was calculated on non-rarefied relative abundance data. Sequences are available at the European Nucleotide Archive (ENA) at EMBL-EBI under project number PRJEB50572, and submission number ERA8814161.

### 2.3. Chemical Analysis

Sulfate, phosphate, and thiosulfate were measured by ion chromatography on a Dionex ICS 6000 equipped with an IonPac AS17-C analytical column (4×2550 mm), and a AS17-C guard column (Dionex, USA) eluted at 30 °C with potassium hydroxide (5 mM, 0.25 mL·min^-1^). Volatile fatty acids (VFAs) were measured with a GC system Agilent 7890B equipped with a flame ionization detector and an HP-FFAP column (25m × 0.32mm). Total organic carbon (TOC) measurements during steady-state were performed using a TIC-TOC analyzer (TOC-L CPH/CPN series, Shimadzu, Japan), equipped with a non-dispersive infrared detector (NDIR). The autosampler settings for inorganic carbon removal required sample acidification with 1 M H_2_SO_4_, flushing with synthetic air (C_x_H_y_<1 ppm, Linde Gas Benelux B.V), and sample injection at 720 °C. Free dissolved sulfide and NH_4_^+^ were analyzed using Hach Lange kits LCK-653 and LCK-303 (Hach, Germany), respectively. Free sulfide samples were diluted in anaerobic water and preserved using a NaOH (12 mM) and zinc acetate solution. Headspace gas composition was analyzed through gas chromatography. Further description of the procedures and equipment follow those by Hidalgo-Ulloa et al. (2020).

## 3. Results

### 3.1. Sulfidogenic productivity

In the neutral pH regime (maintained at 6.9 ± 0.1), steady-state was reached on day 92 of operation, with an average VSPR of 1.787 ± 0.177 g·L^-1^·d^-1^ S^2-^ (Figure 1). In this period, S^0^ reduction accounted for 98.5% of the VSPR, while reduction of SO_4_^2-^, present in the media, accounted for the remaining 1.5 ± 0.5% of the VSPR at steady-state conditions (98 % reduction of the SO_4_^2-^ loaded). In addition, thiosulfate formation was detected in the reactor liquor in this period (Supplementary information, S.I.2).

**Figure 1.**
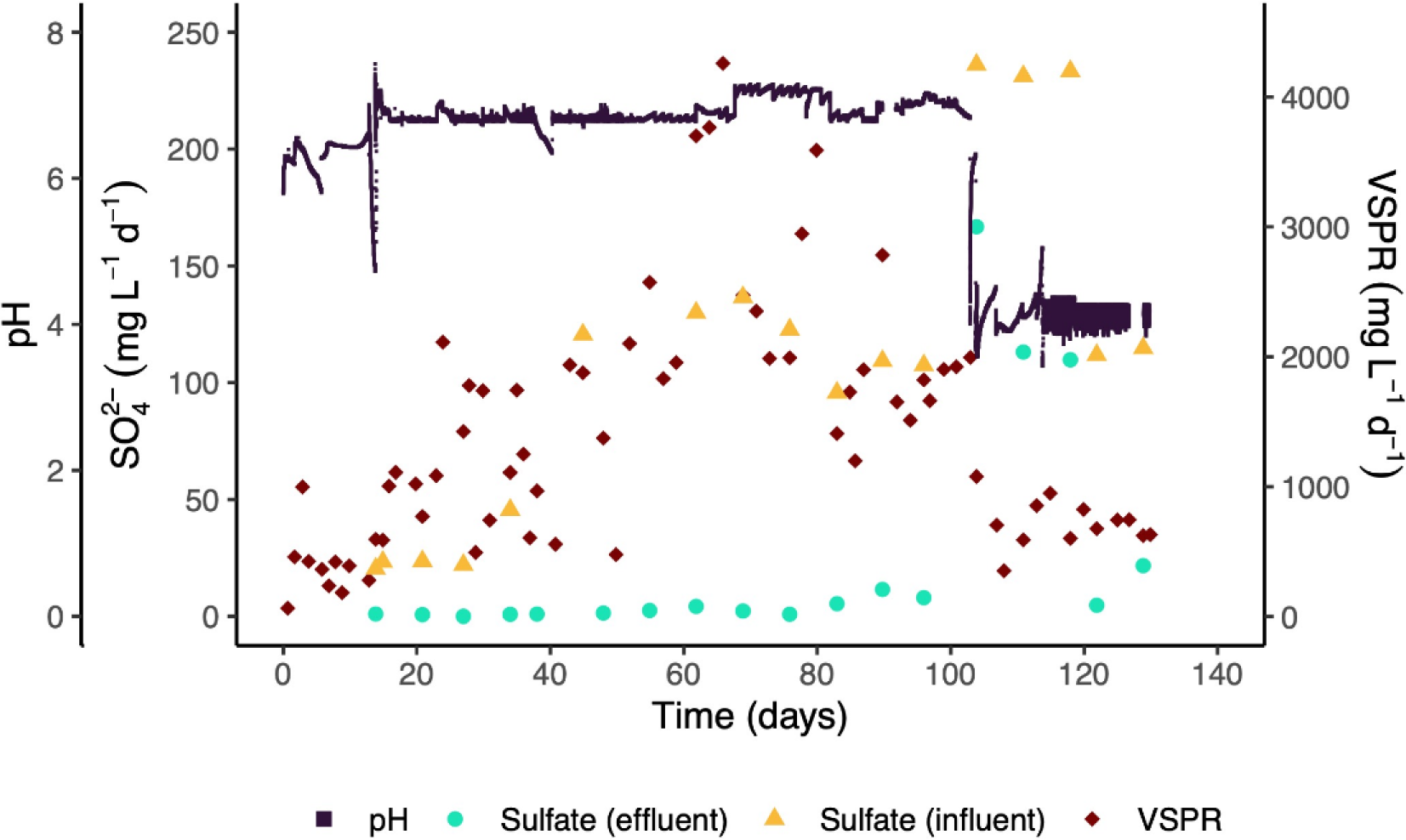
Volumetric sulfide production rates (VSPR) of the Emmen sludge in the 4L gas lift reactor (secondary axis, red markers) contrasted with the pH changes (primary axis, black markers) and SO _4_^2-^ loading rate in the influent (primary axis, yellow markers) and effluent (primary axis, turquoise triangles).

Upon concluding steady-state under neutral pH conditions (days 92-103), the reactor pH was decreased to 3.8 ± 0.1 (day 103) (Figure 1). Steady-state conditions were reached 17 days after the pH decrease (day 120) and sustained through day 130. The VSPR in the steady-state at low pH dropped to 0.705 ± 0.068 g·L^-1^·d^-1^ S^2-^, a 2.5-fold decrease of the VSPR observed during operation at neutral pH. During the initial operation at acidic conditions, the pH was controlled using a 0.1 M H_2_SO_4_ solution. However, on day 118, this solution was replaced with 0.1 M HCl to limit the contribution of SO_4_^2-^ reduction to the VSPR (Figure 1). SO_4_^2-^ reduction accounted for 5.1 ± 0.2% of the total VSPR during this regime. Despite the relative increase of SO_4_^2-^ reduction during operation at acidic pH, the absolute SO_4_ ^2-^ reduction remained equivalent to that at neutral pH. At acidic conditions, the VSPR from SO_4_^2-^ reduction accounted for 0.334 g·L^-1^·d^-1^ S^2-^ while at neutral pH was 0.329 g·L^-1^·d^-1^ S^2-^. No thiosulfate was detected during operation at acidic pH (Supplementary information, S.I.2).

### 3.2. VFA, NH _4_^+^ and biomass concentration changes across pH regimes

During the initial operation in the neutral pH regime, we observed an increase in the VFA concentration, reaching up to 1474 mg·L^-1^ (day 48). Subsequently, VFA concentrations decreased to below the limit of detection (LOD < 2.5 mg·L^-1^) on day 64 (Figure 2). Acetate accounted for 93 ± 5% of total molar VFA (Supplementary information, S.I.3). On day 27 of operation, the NH_4_^+^ concentration in the effluent was below detection (LOD < 0.1 mg·L^-1^ NH_4_^+^) (Figure 2), indicating NH_4_^+^ limitation. This lasted until day 38, when the NH_4_^+^ concentration in the influent was increased. During this period the VSPR fluctuated, reaching a maximum of 4.26 g·L^-1^·d^-1^ S^2-^ (day 66).

**Figure 2.**
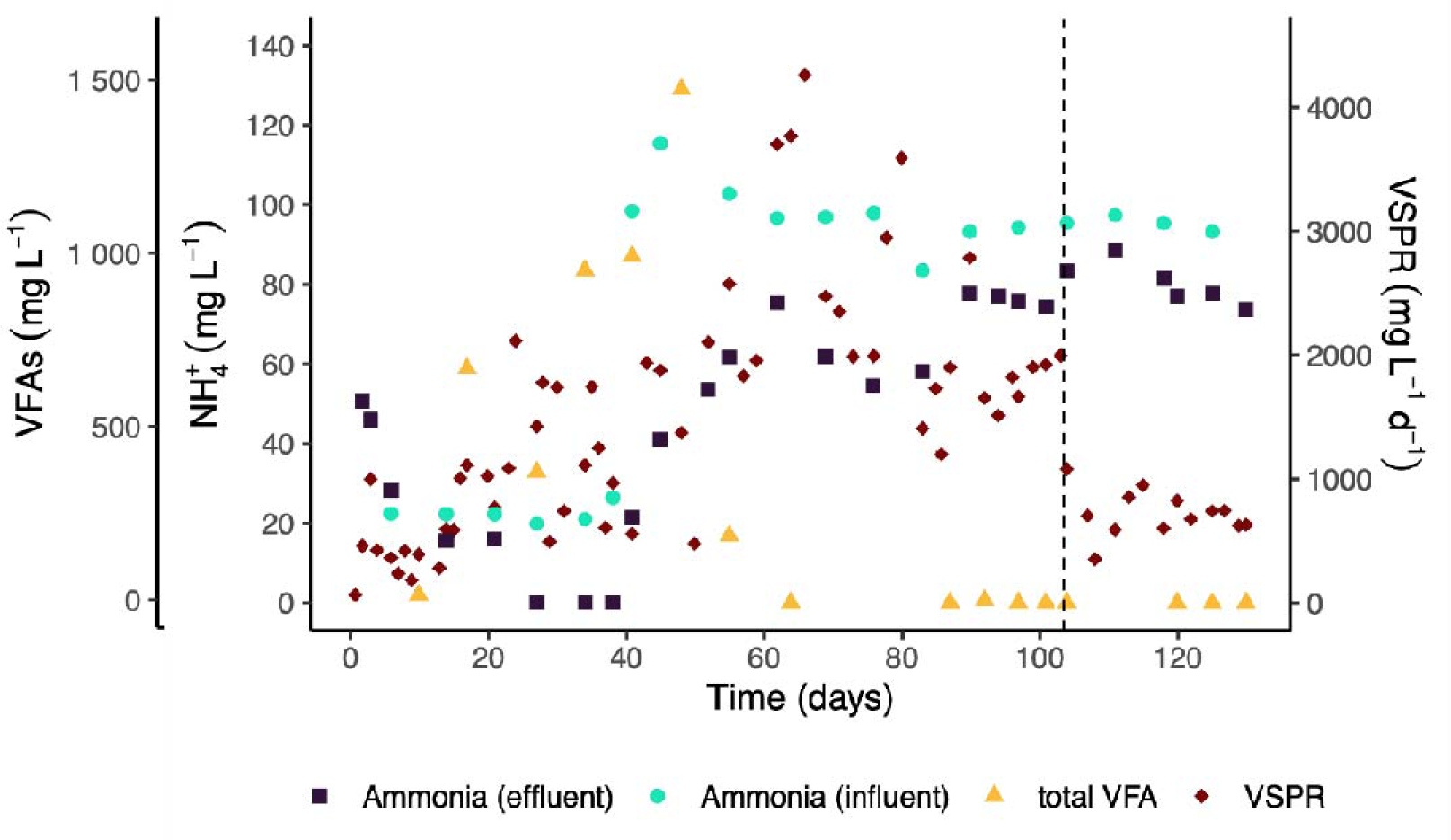
Total volatile fatty acid (VFA) concentration in the reactor (primary axis, yellow markers) and NH _4_^+^ concentrations in the influent (primary axis, black markers) and effluent (primary axis, turquoise markers) and the VSPR (secondary axis, red markers). Dotted line indicates the switch from neutral to acidic pH.

Total organic carbon (TOC) was used as proxy for biomass concentration during the steady-state in both pH regimes, since VFA concentrations were below LOD in these periods. During steady-state at pH 6.9 (day 92 – 103), the TOC concentration was 26.5 ± 1.7 mg·L^-1^, while during steady-state at acidic pH 3.8 this was 18.6 ± 1.0 mg·L^-1^. The TOC concentration was converted into biomass concentrations using the median empirical biomass formula for prokaryotes (CH_1.6_O_0.4_N_0.2_) (Rittmann and McCarty, 2020). This resulted in estimated biomass concentrations of 50.4 ± 3.2 mg·L^-1^ and 35.3 ± 1.8 mg·L^-1^ in the neutral and acidic pH regimes, respectively.

### 3.3. Assessment of H_2_ and CO_2_ flow rates in the VSPR

Possible limitations in mass transfer of the electron (H_2_) and carbon (CO_2_) sources were assessed by increasing H_2_ and CO_2_ influent flow rates. Four different H_2_ flow rate regimes were tested: 5.6 (day 23 – 34), 11.2 (day 34 – 64), 28.0 (day 64 – 80), and 8.0 (day 80 – end) L·h^-1^. Similarly, two CO_2_ gas flow regimes were evaluated 0.7 (day 1 – 80, day 86 – end) L·h^-1^ and 2.0 (day 80 – 86) L·h^-1^. During steady-state operation in both pH regimes, the H_2_ and CO_2_ remained at 8 and 0.7 L·h^-1,^ respectively. Although changes in VSPR were observed upon increasing the H_2_ flow rate on day 64, these changes were not consistent and did not lead to an increased steady-state VSPR. Likewise, increments in the CO_2_ flow rate did not lead to an immediate effect on the VSPR.

**Figure 3.**
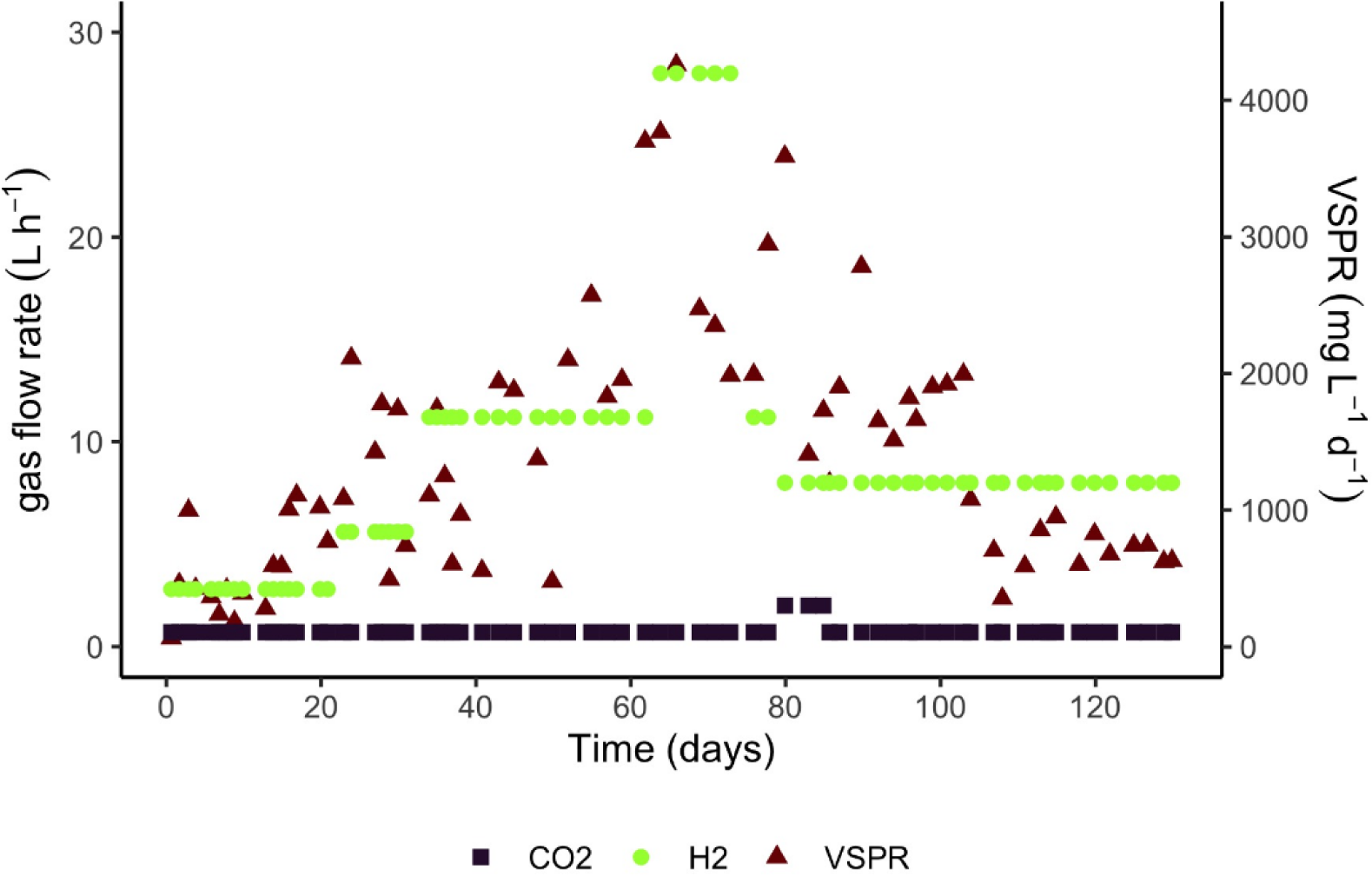
Changes in the influent gas rate (primary axis) and VSPR (secondary axis, red markers). Hydrogen (primary axis, green markers) and carbon dioxide (primary axis, black markers).

### 3.4. Microbial community composition and shifts throughout reactor operation

To assess the effect of the pH decrease on the microbial community composition and determine the dominant (sulfidogenic) microbial taxa in the bioreactor, reactor samples from day 24, 59, 73, and 101 (neutral pH, 6.9) and day 118 and 130 (acidic pH, 3.8) were analyzed with 16S rRNA amplicon sequencing. After filtering and quality control, between 22058 and 696963 reads remained per sample (Supplementary information, S.I.4). The alpha diversity decreased upon lowering the pH: at pH 3.8, the dominance was 0.69 ± 0.05, compared to 0.42 ± 0.07 at pH 6.9 (Figure 4A). For the inoculum and the S^0^ a dominance of 0.23 ± 0.03 and 0.28 ± 0.03 was calculated, respectively. Comparison of the beta diversity indicated a clear separation between reactor samples from the two pH regimes, the inoculum, and the S^0^ (Figure 4B). Even though steady-state was reached only on day 92, the beta diversities on day 24 and day 101 were already highly similar (Figure 4B).

**Figure 4:**
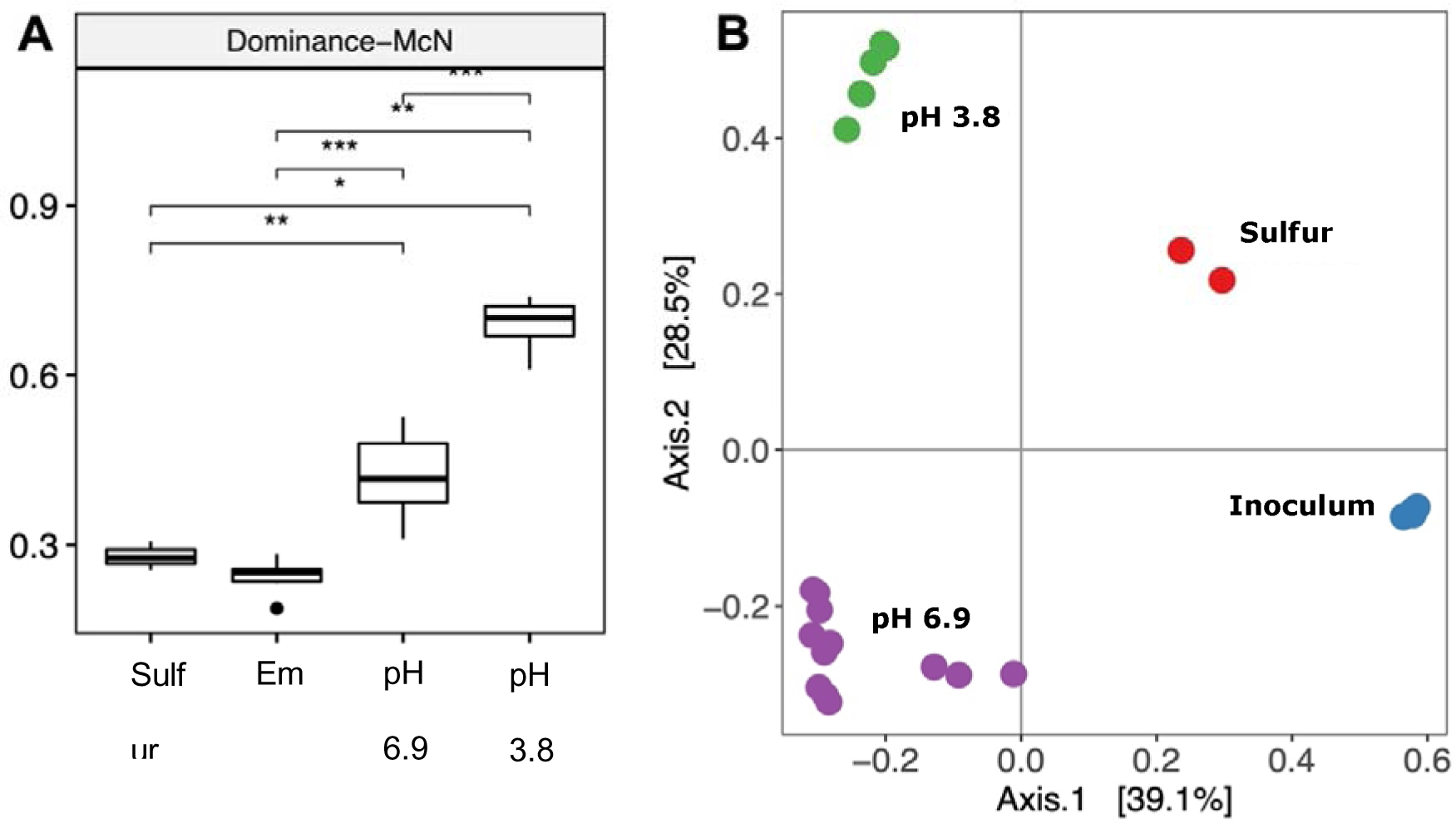
(A) Alpha diversity of samples from S^0^, inoculum (Emm), and the S _8_^0^-reducing reactor at pH 6.9 and pH 3.8 expressed as McNaughtons Dominance. Statistical significance of the difference between means of the four groups was determined using the Wilcoxon rank -sum test. Significance is indicated by “*” where “****”, “***”, “**”, and “*”, correspond to a p-value less than 0.0001, 0.001, 0.01, 0.05, and and “n.s” indicates the difference is not significant. (B) Principal Coordinates Analysis (PCoA) plot comparing the beta diversities (Bray-Curtis dissimilarity index) between samples from S^0^ (red), inoculum (Emm, blue), and the S_8_^0^-reducing reactor at pH 6.9 (purple) and pH 3.8 (green).

Different taxa were detected at pH 6.9 compared to pH 3.8 (Figure 5), and the dominant taxa in samples from both regimes differed from the original inoculum and the added S (Supplementary information, S.I.5). Of the ten most abundant taxa detected at both pH regimes (Figure 5), *Sulfurospirillum, Sulfurovum, Desulfovibrio, Acetobacterium*, and an unknown genus from the order OPB41 within the *Coriobacteria* class (*Actinobacteria* phylum) were abundant at pH 6.9, but decreased to below detection at pH 3.8. Conversely, *Thiomonas* and *Thermodesulfobium* were abundant at pH 3.8, but not detectable (*Thermodesulfobium*) or only present at 1.0 ± 0.2 % (*Thiomonas)* during operation at pH 6.9. Reads classified as *Desulfurella*, *Methanobacterium*, and *Microbacter*, were present throughout both pH regimes, with *Desulfurella* becoming highly dominant at pH 3.8.

**Figure 5:**
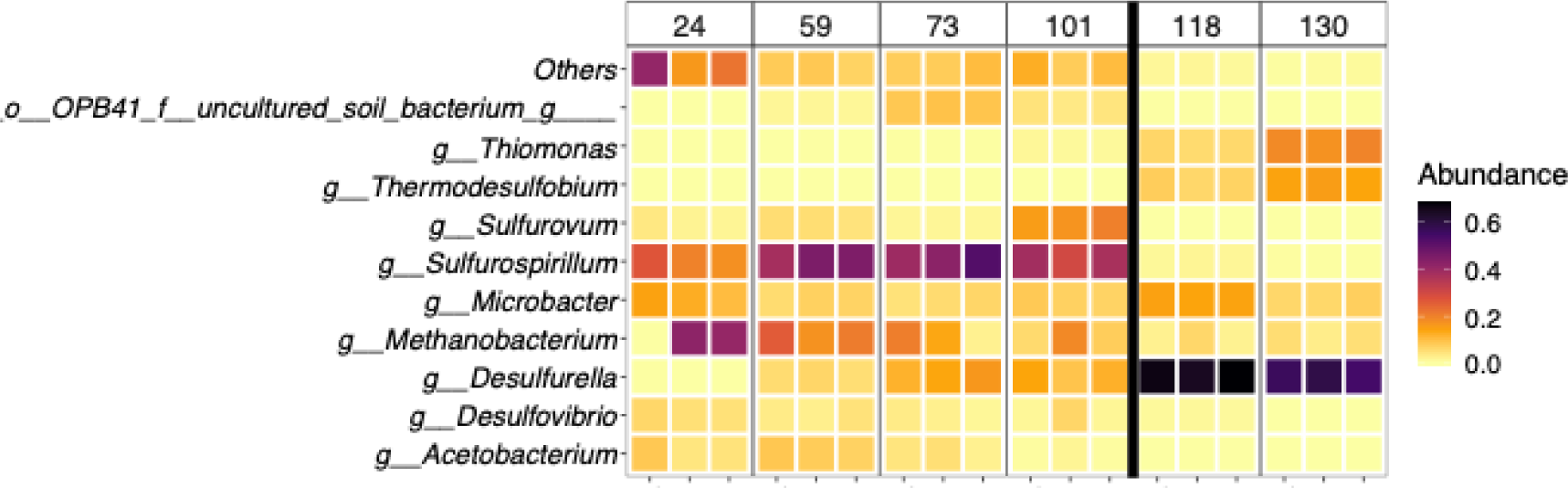
Top 10 most abundant taxa according to sequenced reads in samples from the neutrophilic (pH 6.9) and acidophilic (pH 3.8) operating regimes, with remainder grouped under ’others’. Colors represent the relative abundance of sequenced reads assigned to the taxa indicated on the y-axis. Individual replicates from triplicate samples are shown, grouped per day of sampling. The black line indicates the separation of the neutrophilic (days 24 – 101) and acidophilic (day 118 & 130) regimes. o_—_: order; g_—_: genus.

During operation at pH 6.9, reads assigned to the genus *Sulfurospirillum* were most abundant, increasing from 21.2 ± 5.0 % on day 24 to 44.4 ± 6.7 % on day 73, then slightly decreasing to 35.0 ± 4.7 % on day 101 before dropping to 0.25 ± 0.1 % upon the transition to pH 3.8. Similarly, reads assigned to the genus *Sulfurovum* increased in relative abundance during operation at pH 6.9, from 2.3 ± 1.1 % on day 24 to 17.1 ± 2.5 % on day 101, but dropped to below LOD upon the decrease in pH. The relative abundance of *Desulfovibrio* and *Acetobacterium* decreased throughout the neutrophilic regime, from 5.1 ± 0.8 % and 5.5 ± 2.5 % on day 24 to 2.9 ± 2.8 % and 0.8 ± 0.7 % on day 101, respectively, and dropped below LOD at acidic pH. Sequences classified as *Thermodesulfobium* were not detected at pH 6.9, and sequences related to *Thiomonas* were detected only at low abundance, between 0.04 ± 0.08 % on day 24 to 1.0 ± 0.2 % on day 101. However, upon the transition to pH 3.8, reads assigned to *Thermodesulfobium* and *Thiomonas* increased to 14.2 ± 0.7 % and 8.2 ± 1.2 %, respectively, by day 130, Of the three top ten taxa present in both pH regimes, *Methanobacterium and Microbacter* were already detected on day 24. The abundance of *Methanobacterium* decreased during operation at pH 6.9, from 27.5 ± 23.7 % on day 24 to 10.5 ± 6.9 % on day 101, whereas the abundance of *Microbacter* remained approximately constant, between 12.2 ± 2.0 % on day 24 and 7.2 ± 0.8 % on day 101. *Desulfurella*, also detected at both pH values, accounted for < 0.1 % of reads on day 24, and then increased to 11.5 ± 2.4 % on day 101. Upon the switch to acidic pH, *Desulfurella* became the dominant taxon in the sequenced reads, accounting for 66.4 ± 1.8 % of total sequenced reads on day 118 and 56.3 ± 2.3 % on day 130.

## 4. Discussion

### 4.1. Autotrophic and heterotrophic S^0^ reduction at neutral pH

During operation at neutral pH, the VSPR exhibited a sinusoidal pattern before reaching steady-state. The changes observed in the VFA concentrations, predominantly acetate, suggest that these could be driving the fluctuations in VSPR. Prior to reaching steady-state, acetate concentrations increased up to 1474 mg·L^-1^ by day 48, and then decreased, coinciding with a peak in the VSPR (4.260 g S^2-^·L^-1^·d^-1^) by day 66.

The link between fluctuations in VSPR and VFA oxidation appears to be supported by the observed changes in the NH_4_^+^ consumption rates. On day 27, the NH_4_^+^ concentration in the effluent was below LOD (0.1 mg·L^-1^), indicating it was potentially limiting microbial growth. The subsequent increase in NH_4_^+^ concentration in the influent (day 38) led to a surge in the NH_4_^+^ consumption rate from 11 mg NH_4_^+^·L^-1^·d^-1^ (day 38) to 31 mg NH_4_^+^·L^-1^·d^-1^ (days 41-45), suggesting an increase in microbial biomass. The mass balance analysis over the metabolite production, presuming H_2_ as the sole electron donor, indicated that the observed increase in biomass was primarily driven by acetogenesis. Specifically, up to 59 % of the electron donor consumption was allocated for acetate formation (day 41), highlighting the competitive dynamics between acetogenesis and sulfidogenesis under these conditions.

This sinusoidal pattern continued until day 80, with fluctuations in the NH_4_^+^ consumption rates followed by a transient spike in the VSPR and decreasing concentrations of VFA’s. However, the amplitude and frequency of the fluctuations progressively decreased, until reaching steady-state with stable VSPR and NH_4_^+^ consumption rates (1.78 ± 0.17 g S^2-^·L^-1^·d^-1^; 6.95 ± 0.65 mg NH_4_^+^·L^-1^·d^-1^) and VFA concentrations below LOD. The simultaneous decrease in acetate concentration and rate of NH_4_^+^ consumption in the reactor compared to the values observed before reaching steady state suggests a decreased activity of acetogenic microorganisms.

The detection of acetate and its apparent consumption indicates the occurrence of heterotrophic S^0^ reduction. This is supported by the microbial community composition observed in this period. *Acetobacterium*, with the type strain *A. woodii*, is a well-studied acetogen, capable of acetate production from H_2_ and CO_2_ (Balch et al., 1977). The dominant S^0^-reducing taxa detected in the sequenced reads during operation at pH 6.9, support the occurrence of both heterotrophic and autotrophic S^0^ reduction. *Sulfurospirillum* was the dominant S^0^-reducing taxon, followed by *Sulfurovum*, *Desulfurella*, and *Desulfovibrio.* While *Sulfurovum*, *Desulfurella*, and *Desulfovibrio* species are capable of autotrophic S^0^ reduction, *Sulfurospirillum* species described to date, such as *S. arcachonense* (Finster et al., 1997; Stolz et al., 1999) *S. deleyianum* (Schumacher et al., 1992; Wolfe and Pfennig, 1977), and *S. Diekertiae* (Jin et al., 2023) are capable of S^0^ reduction with H_2_ as electron donor but cannot use CO_2_ as carbon source, requiring organic compounds such as acetate and supporting the hypothesis that the VFA were used as organic carbon source (Figure 6). Previous studies of neutrophilic S^0^-reducing bioreactors fed with glucose and acetate also detected *Sulfurospirillum* as one of the dominant S^0^-reducing taxa (Qiu et al., 2017). So far, no acidophilic *Sulfurospirillum* species have been described, explaining their disappearance during operation at acidic pH. While *Sulfurospirillum* was the dominant genus between day 24 and day 73, steady-state was only reached on day 92, with a VSPR lower than observed in the period before steady state. At steady-state on day 101, *Sulfurospirillum* remained a dominant community member, with an increased abundance of *Sulfurovum*. Several *Sulfurovum* species are capable of autotrophic S^0^ reduction with H and CO, e.g. *Sulfurovum aggregans* and *Sulfurovum* sp. NBC37-1 (Mino et al., 2014) (Nakagawa et al., 2005; Yamamoto et al., 2010). Together with *Desulfurella* these likely accounted for autotropic S^0^ reduction during steady state. The sharp decrease in NH_4_^+^ consumption and in the relative abundance of reads assigned to *Acetobacterium* during steady-state could indicate that the availability of organic carbon limited growth of *Sulfurospirillum*. The decrease of *Acetobacterium* could be related to increasing H_2_S concentrations, as was observed in homoacetogenic mixed cultures at concentrations above 3.3 mM (Ntagia et al., 2020). In summary, while heterotrophic S^0^ reduction was dominant in the first 80 days, autotrophic S^0^ reduction appeared to be dominant during steady-state.

**Figure 6:**
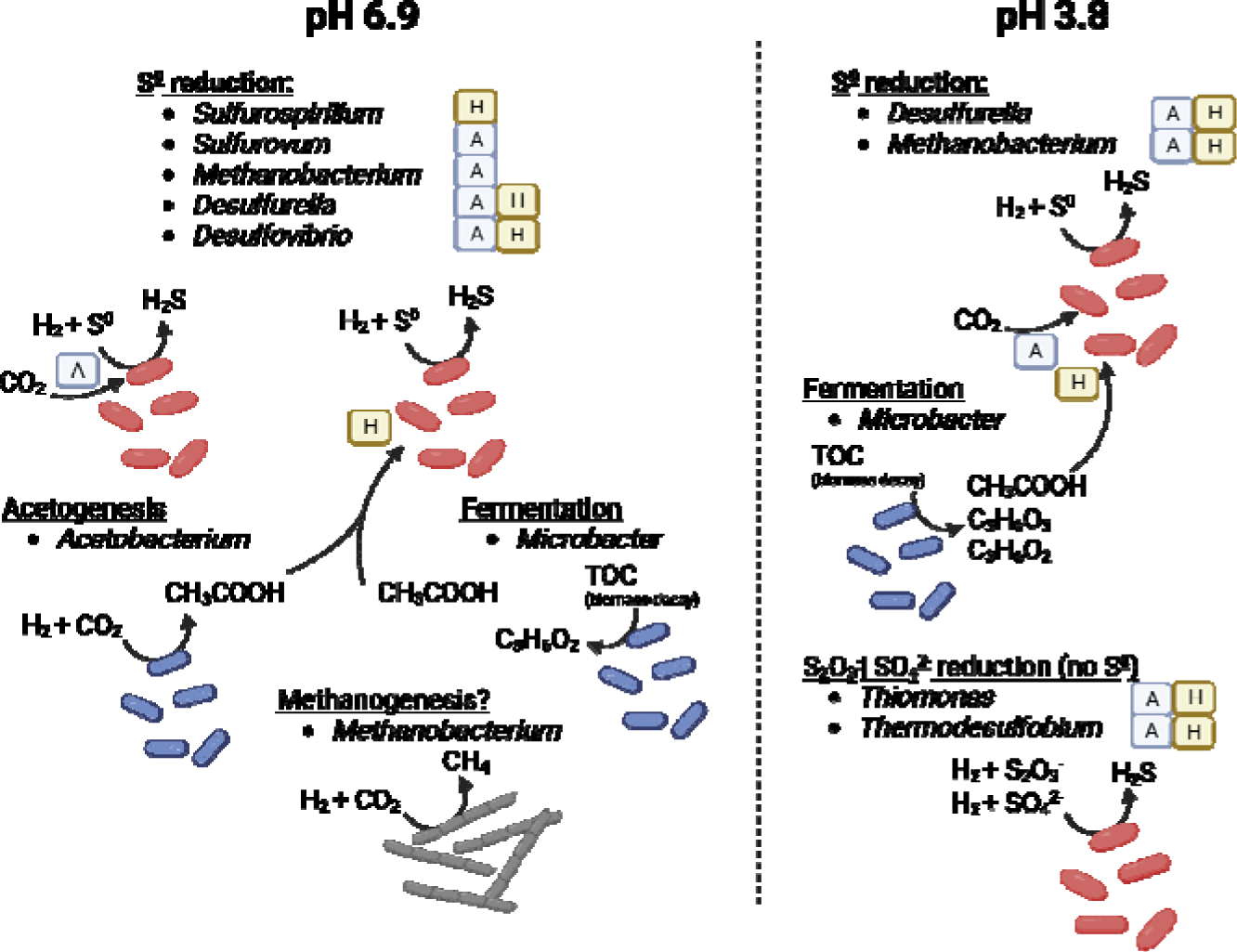
Schematic representation of proposed dominant metabolic reactions and associated microbial taxa detected in the 16S rRNA gene amplicon sequencing reads obtained for the pH 6.9 and pH 3.8 regime. A: autotrophic, H: heterotrophic.

### 4.2. CO _2_ and H _2_ limitations during circumneutral pH operation

We found no correlation between the VSPR and the influent gas flow rates. In the interval from days 41 to 52, during the first H_2_ flow rates increase, previously adjusted to 11.2 L^-1^·d^-1^on day 34, the VSPR remained similar to those observed under lower H_2_ inflow conditions (days 23 - 34). This finding suggests that the relationship between H_2_ flow rates and VSPR might be more complex and that other factors, such as the dynamics between VFA production and oxidation discussed above, influenced the VSPR.

From days 55 to 64, despite the absence of further modifications to the H_2_ flow, the VSPR increased, to 3.8 g S^2-^·L^-1^·d^-1^ by day 64. This trend continued after the increase in H_2_ flow to 28 L·h^-1^ initiated on day 64, increasing the VSPR to a peak of 4.2 g S^2-^·L^-1^·d^-1^ by day 66. This peak, rather than being a direct consequence of the increase in H_2_ flow rate, appears to be a continuation of increasing trend noted earlier, suggesting that VSPR dynamics are influenced by more factors than H_2_ supply alone.

The subsequent decline in VSPR post-peak, despite the sustained increase in H_2_ flow, could suggest carbon limitation, potentially due to the unadjusted CO_2_ flow rates. An increase in H_2_ inflow can dilute CO_2_ partial pressure, potentially impacting both autotrophic S^0^ reduction and the production of VFAs necessary for heterotrophic S^0^ reduction. This potential limitation is challenged, however, by the observed NH_4_^+^ consumption rates, which serve as indicators of microbial growth. Specifically, NH_4_^+^ consumption rates increased from 8 mg NH_4_^+^·L^-1^·d^-1^ on day 62 to 15 mg NH_4_^+^·L^-1^·d^-1^ by day 69, indicating an increase in microbial activity and an increased requirement for CO_2_ fixation for new biomass production, thereby suggesting that CO_2_ was not a limiting factor during this timeframe. Moreover, a subsequent increase in CO_2_inflow rates between days 80 and 86 did not notably influence the VSPR, further supporting that gas inflow rates were not the primary drivers of the observed changes in VSPR. Taken together, this underscores the need to consider the intricate interplay among gas flows, microbial metabolism, and environmental conditions when analyzing the factors influencing VSPR in sulfidogenic bioreactors.

### 4.3. Shifting of the microbial community upon acidification

A clear change in microbial community diversity (Figure 4) and composition (Figure 5) was observed in the reactor samples upon the switch from neutral (pH 6.9, days 0 - 101) to acidic conditions (pH 3.8, days 101 - 130). Figure 5 The absence of *Sulfurospirillum* and *Sulfurovum* from the sequenced reads after the shift to acidic pH is in line with the reported neutrophilic physiology of *Sulfurospirillum* (Finster et al., 1997; Stolz et al., 1999) (Schumacher et al., 1992; Wolfe and Pfennig, 1977) and *Sulfurovum* species (Inagaki et al., 2004), (Mino et al., 2014) (Nakagawa et al., 2005; Yamamoto et al., 2010). Upon the switch to acidic pH, *Desulfurella* became the dominant S^0^-reducing genus according to the sequenced reads. *Desulfurella* was already one of the dominant S^0^-reducing taxa during operation at pH 6.9, suggesting an important role for this genus throughout reactor operation. Closer inspection of the individual ASVs classified as *Desulfurella* showed that 31 different ASVs were detected throughout reactor operation, of which 13 occurred in samples from both pH 6.9 and pH 3.8, 2 were unique to samples from pH 6.9, and 16 were unique to pH 3.8. Although 1 ASV consistently accounted for 77 to 100 % of reads classified as *Desulfurella* in all samples, the variation among the other ASVs could indicate that another, more acidophilic *Desulfurella* species became dominant at pH 3.8. All *Desulfurella* species, both neutrophilic and acidophilic, are capable of S^0^ reduction, with *D. acetivorans* (Bonch-Osmolovskaya et al., 1990)*, D. kamchatkensis* and *D. propionica* (Miroshnichenko et al., 1998), and *D. multipotens* (Miroshnichenko et al., 1994) growing between pH 6.7 and 7.2, and *D. amilsii* at pH 3.8 – 6.9 (Florentino et al., 2016a). *Desulfurella* species can utilize both organic and inorganic substrates as carbon and energy sources. This versatility is reflected by the dominance of *Desulfurella* in experiments performed under different conditions, such as acidophilic S^0^-reducing enrichments using either acetic acid, methanol, or H_2_/CO_2_ as energy and carbon source (Florentino et al., 2015) and in heterotrophic (Guo et al., 2021; Guo et al., 2019) and autotrophic reactors (current study) at neutral and acidic pH.

Next to *Desulfurella*, reads assigned to *Thiomonas* and *Thermodesulfobium* increased in relative abundance at pH 3.8. *Thiomonas* was already observed at low abundance on the final day of sampling at neutral pH, but *Thermodesulfobium* remained below detection before the switch to acidic pH. *Thiomonas* species have been isolated predominantly from acid mine drainage sediments and hot spring environments (Akob et al., 2020), and grow at acidic pH, with minimum pH 3.0 for *T. metallidurans* (Akob et al., 2020), to neutral pH, with a maximum pH of growth of 8.5 for *T. bhubaneswarensis)* (Panda et al., 2009). Furthermore, *T. islandica* is capable of autotrophic growth on H_2_, and utilization of organic and inorganic energy and carbon sources (Vésteinsdóttir et al., 2011). However, chemolithotrophic growth with H_2_ was not confirmed with S^0^, raising the question of which energy metabolism is utilized by the *Thiomonas* species detected in the reactor. Like *Thiomonas*, *Thermodesulfobium* species were reported to grow chemolithoautotrophically with H_2_/CO_2_, using oxidized sulfur compounds such as SO_4_^2-^ or thiosulfate as electron acceptors, but were not able to use S^0^ (Frolov et al., 2017; Mori et al., 2003). Its potential role as SO_4_^2-^ reducer could be further supported by the observation that SO_4_^2-^ concentrations again started decreasing 19 days after the pH decrease (Figure 1).

The high relative abundance of reads assigned to *Methanobacterium* at both neutral and acidic conditions could indicate the occurrence of methanogenesis from H_2_ and CO_2_. During operation at pH 6.9, methane was not monitored, and its formation in the first 103 days can therefore not be excluded. During operation at pH 3.8, the gas composition was measured on day 104 and day 129, but no CH_4_ was detected. We propose that rather than methanogenesis, *Methanobacterium* performed S^0^ reduction with H_2_ as electron donor, as demonstrated previously (Stetter and Gaag, 1983). More recently, *Methanobacterium* was implicated as the S^0^ reducing species responsible for unwanted H_2_S formation from S^0^ formed in an H_2_S removal process (Zhou et al., 2011).

### 4.4. Implications of the acidification for the VSPR

As previously discussed, upon the transition from pH 6.9 to pH 3.8, a 2.3-fold decrease in the VSPR at steady-state was observed, accompanied by a change in relative microbial community composition and a reduction of diversity. The decrease in relative abundance of reads assigned to the inferred S^0^-reducing genera *Sulfurospirillum, Desulfurella, Sulfurovum, Desulfovibrio,* and *Methanobacterium* from 77.9 % on neutral pH steady-state (day 101), to 60.1 % on the acidic steady-state (day 130) could partly explain the decrease in VSPR. However, no absolute abundance data was obtained, serving this finding as an indication. Nevertheless, the VSPR changes are likely the result of a larger multifactorial effect, as further discussed below.

TOC measurements indicated a decrease in estimated biomass concentrations from 50.4 ± 3.2 mg·L^-1^ at the steady-state at neutral pH to 35.3 ± 1.8 mg·L during the steady-state at pH 3.8. Interestingly, the rates of NH_4_^+^consumption were similar in both steady-states (6.95 ± 0.65 mg NH_4_ ·L^-1^·d^-1^, days 92 – 103; 6.97 ± 0.78 mg NH_4_^+^·L^-1^·d^-1^, days 120 – 130). The decrease in biomass concentration, together with a decrease in VSPR but a similar NH_4_ ^+^ consumption rate upon the acidification, can likely be explained by the increased cellular maintenance energy requirements for acid stress resistance mechanisms such as active proton export and selective membrane permeability required for survival at low pH (Hu et al., 2020) (Guan and Liu, 2020).

Furthermore, it has been hypothesized that microbial S^0^ conversion rates are limited due to the low S^0^ solubility in water (53.8 nM, 28 °C) (Florentino et al., 2015) rendering it predominantly solid in water (Kamyshny, 2009). Because polymeric sulfur chains exhibit a bonding energy that is 2.4 kJ·mol^-1^ lower than cyclooctasulfur bonds (Franz et al., 2007), it has been suggested that polysulfides (S_n_ ^2-^) are the primary terminal electron acceptor in S^0-^-reducing processes (Hedderich et al., 1998; Schauder and Müller, 1993). However, S_n_ ^2-^ chain length is limited under acidic conditions resulting in a low polysulfide concentration. While, at neutral pH, the average S_n_ ^2-^ chain length is four to six sulfur atoms (S_4_ ^2-^ - S_6_ ^2-^), at acidic conditions this is limited to two sulfur atoms (S-S^2-^) (Kamyshny et al., 2007). Moreover, H_2_S_n_ might be the dominant polysulfide form under acidic conditions, which has been reported to be almost insoluble in water (Steudel, 2020). Hence, it remains to be determined whether the limitation in microbial S^0^ –reduction at acidic pH is due to the bioavailability of S^0^ forms utilized by acidophilic S^0^-reducers or by limitations in the S^0^ transfer rate (Boyd and Druschel, 2013; Florentino et al., 2016a; Takahashi et al., 2010).

By using the same reactor set-up and anaerobic sludge as inoculum as in our prior work (Hidalgo-Ulloa et al., 2023), we aimed to determine the extent to which S^0^ availability influenced the VSPR. The VSPR in this study showed a 2.6-fold increase under the acidic conditions compared to that reported under equivalent pH conditions but at higher temperature (0.27 g S^2-^·L^-1^·d^-1^ at 60 °C) (Hidalgo-Ulloa et al., 2023). These differences appear despite the solubility of S^0^ increases by one order of magnitude at 60 °C relative to the temperature in the current study (Kamyshny, 2009), suggesting increased bioavailability and thereby enabling higher rates. However, the industrial sludge used as inoculum originated from a process operated at mesophilic conditions. Therefore, the S^0^ conversion limitations under thermoacidophilic conditions were likely the result of microbial growth limitations rather than limitations on S^0^ availability.

Nevertheless, the VSPR in this study aligns closely with those documented under similar conditions. For instance, Guo et al. (2021) reports VSPR reaching 0.844 ± 0.115 g S^2-^·L^-1^·d^-1^at pH 3.8 and 25 °C while Sun et al. (2020b) found VSPR up to 0.845 g S^2-^·L^-1^·d^-1^ at pH 3.5 and room temperature (not specified). Furthermore, the VSPR at the circumneutral pH is also in the same order of magnitude of those reported from research performed at laboratory-scale (Escobar et al., 2007; Zhang et al., 2018a) and industrial applications (Gonzalez-Contreras et al., 2016), under comparable pH and temperature conditions. This consistency in VSPR across studies, transcending differences in inoculum, electron donor, S^0^ source, operational parameters, reactor configurations, and scale, hints at an intrinsic limit in microbial S^0^-reduction rates. Specifically, under acidic conditions (pH < 4), the VSPR from S^0^-reduction appears to plateau in a 10^-1^ g S^2-^·L^-1^·d^-1^ order of magnitude, while at neutrophilic conditions (pH 6.8 – 7.5), this seems to extend to a 10^0^ g S^2-^·L^-1^·d^-1^ order of magnitude.

## Conclusions

The findings of this study, along with our previous reports of this inoculum at high temperature and acidic conditions, suggest that temperature has a more pronounced effect on the VSPR than pH when using the Emmen granular sludge as seed. While at mesophilic temperatures the VSPR dropped 2.5-fold when the pH was decreased from 6.9 to 3.8, this was still around 2.6-fold higher than the VSPR observed at thermoacidophilic conditions (pH 3.5, 60 °C). Although the differences in VSPR are likely related to the original growth conditions of the inoculum, the VSPR at acidic conditions corresponds well with findings from other studies, suggesting a limit on the microbial sulfidogenic rate under acidic conditions (pH < 4).

During the initial operation stages at neutral pH, both autotrophic and heterotrophic sulfidogenesis occurred, despite the chemoautotrophic operating conditions. Upon the switch to acidic conditions, the microbial community became dominated by *Desulfurella.* Analysis of individual ASV assigned to this taxon suggests the presence of different species from that at pH 6.9.

## Supporting information

Supplementary material 1 to 5, described in the text.

## Abbreviations

HRT: hydraulic retention time
LOD: limit of detection
S^0^: elemental sulfur
TOC: total organic carbon
VFA: volatile fatty acids
VSPR: volumetric sulfide production rates.

## Footnotes

* Change in sulfide concentration 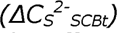 in the effluent scrubber over time, V _SCB_ is the scrubber volume (1.8 L), V _R_ is the effective working reactor volume (4 L), and t _n_ and t_n+1_ are the sampling time (day) at the initial time (n) and final sampling time (n+1), respectively.

† Steady-state was considered reached when the standard deviation of the average VSPR remained within a 10% deviation during ten consecutive operational days (4x HRT).

## References

Adams, M., Lawrence, R. and Bratty, M. 2008. Biogenic sulphide for cyanide recycle and copper recovery in gold–copper ore processing. Minerals Engineering 21(6), 509–517.

Akob, D.M., Hallenbeck, M., Beulig, F., Fabisch, M., Küsel, K., Keffer, J.L., Woyke, T., Shapiro, N., Lapidus, A., Klenk, H.-p. and Chan, C.S. 2020. Mixotrophic Iron-Oxidizing Thiomonas Isolates from an Acid Mine Drainage-Affected Creek. Applied and Environmental Microbiology 86(24).

Azabou, S., Mechichi, T. and Sayadi, S. 2007. Zinc precipitation by heavy-metal tolerant sulfate-reducing bacteria enriched on phosphogypsum as a sulfate source. Minerals Engineering 20(2), 173–178.

Balch, W.E., Schoberth, S., Tanner, R.S. and Wolfe, R.S. 1977. Acetobacterium, a new genus of hydrogen-oxidizing, carbon dioxide-reducing, anaerobic bacteria. International Journal of Systematic Bacteriology 27(4), 355–361.

Batty, J.D. and Rorke, G.V. 2006. Development and commercial demonstration of the BioCOP™thermophile process. Hydrometallurgy 83(1-4), 83–89.

Biebl, H. and Pfennig, N. 1977. Growth of sulfate-reducing bacteria with sulfur as electron acceptor. Archives of Microbiology 112(1), 115–117.

Bonch-Osmolovskaya, E.A., Sokolova, T.G., Kostrikina, N.A. and Zavarzin, G.A. 1990. Desulfurella acetivorans gen. nov. and sp. nov. —a new thermophilic sulfur-reducing eubacterium. Archives of Microbiology 153(2), 151–155.

Boyd, E.S. and Druschel, G.K. 2013. Involvement of intermediate sulfur species in biological reduction of elemental sulfur under acidic, hydrothermal conditions. Applied and Environmental Microbiology 79(6), 2061–2068.

Core Team, R. 2021 R: A lanuguage and environment for statistical computing, R Foundation for Statistical Computing, Vienna.

Escobar, C., Bravo, L., Hernández, J. and Herrera, L. 2007. Hydrogen sulfide production from elemental sulfur by Desulfovibrio desulfuricans in an anaerobic bioreactor. Biotechnology and Bioengineering 98(3), 569–577.

Finster, K., Liesack, W. and Tindall, B.J. 1997. Sulfurospirillum arcachonense sp. nov., a new microaerophilic sulfur-reducing bacterium. International Journal of Systematic Bacteriology 47(4), 1212–1217.

Florentino, A.P., Brienza, C., Stams, A.J.M. and Sánchez-Andrea, I. 2016a. Desulfurella amilsii sp. nov., a novel acidotolerant sulfur-respiring bacterium isolated from acidic river sediments. International Journal of Systematic and Evolutionary Microbiology.

Florentino, A.P., Weijma, J., Stams, A.J.M. and Sánchez-Andrea, I. 2015. Sulfur reduction in acid rock drainage environments. Environmental Science & Technology 49(19), 11746–11755.

Florentino, A.P., Weijma, J., Stams, A.J.M. and Sánchez-Andrea, I. (2016b), pp. 141–175, Springer International Publishing Switzerland.

Franz, B., Lichtenberg, H., Hormes, J., Modrow, H., Dahl, C. and Prange, A. 2007. Utilization of solid ’elemental’ sulfur by the phototrophic purple sulfur bacterium Allochromatium vinosum: A sulfur K-edge X-ray absorption spectroscopy study. Microbiology 153(4), 1268­1274.

Frolov, E.N., Kublanov, I.V., Toshchakov, S.V., Samarov, N.I., Novikov, A.A., Lebedinsky, A.V., Bonch-Osmolovskaya, E.A. and Chernyh, N.A. 2017. Thermodesulfobium acidiphilum sp. nov., a thermoacidophilic, sulfate-reducing, chemoautotrophic bacterium from a thermal site. International Journal of Systematic and Evolutionary Microbiology 67(5), 1482–1485.

Gonzalez-Contreras, P.A., Dijkman, H., Choi, Y. and Espinel, R. 2016 Increasing output of metals at the Pueblo Viejo mine by precipitating copper sulfide using biological H2S generation, Kobe, JP.

Guan, N. and Liu, L. 2020. Microbial response to acid stress: mechanisms and applications. Applied Microbiology and Biotechnology 104(1), 51–65.

Guo, J., Li, Y., Sun, J., Sun, R., Zhou, S., Duan, J., Feng, W., Liu, G. and Jiang, F. 2021. pH-dependent biological sulfidogenic processes for metal-laden wastewater treatment: Sulfate reduction or sulfur reduction? Water Research 204(August), 117628–117628.

Guo, J., Wang, J., Qiu, Y., Sun, J. and Jiang, F. 2019. Realizing a high-rate sulfidogenic reactor driven by sulfur-reducing bacteria with organic substrate dosage minimization and cost-effectiveness maximization. Chemosphere 236, 124381–124381.

Hedderich, R., Klimmek, O., Kröger, A., Dirmeier, R., Keller, M. and Stetter, K.O. 1998. Anaerobic respiration with elemental sulfur and with disulfides. FEMS Microbiology Reviews 22(5), 353–381.

Hedrich, S., Joulian, C., Graupner, T., Schippers, A. and Guézennec, A.G. 2018. Enhanced chalcopyrite dissolution in stirred tank reactors by temperature increase during bioleaching. Hydrometallurgy 179, 125–131.

Hidalgo-Ulloa, A., Buisman, C. and Weijma, J. 2022. Metal sulfide precipitation mediated by an elemental sulfur-reducing thermoacidophilic microbial culture from a full-scale anaerobic reactor. Hydrometallurgy 213, 105950.

Hidalgo-Ulloa, A., Sánchez-Andrea, I., Buisman, C. and Weijma, J. 2020. Sulfur Reduction at Hyperthermoacidophilic Conditions with Mesophilic Anaerobic Sludge as the Inoculum. Environmental Science and Technology 54(22), 14656–14663.

Hidalgo-Ulloa, A., van der Graaf, C.M., Sanchez-Andrea, I. and Buisman, C.J.N. 2023. Continuous single-stage elemental sulfur reduction and copper sulfide precipitation under thermoacidophilic conditions. Water Research, 119948.

Hu, W., Feng, S., Tong, Y., Zhang, H. and Yang, H. 2020. Adaptive defensive mechanism of bioleaching microorganisms under extremely environmental acid stress: advances and perspectives. Biotechnology Advances 42, 107580.

Huisman, J.L., Schouten, G. and Schultz, C. 2006. Biologically produced sulphide for purification of process streams, effluent treatment and recovery of metals in the metal and mining industry. Hydrometallurgy 83(1-4), 106–113.

Hulshoff, L.W., Lens, P.N.L., Weijma, J. and Stams, A.J.M. 2001. New developments in reactor and process technology for sulfate reduction. Water Science and Technology 44(8), 67–76.

Inagaki, F., Takai, K., Nealson, K.H. and Horikoshi, K. 2004. Sulfurovum lithotrophicum gen. nov., sp. nov., a novel sulfur-oxidizing chemolithoautotroph within the ε-Proteobacteria isolated from Okinawa trough hydrothermal sediments. International Journal of Systematic and Evolutionary Microbiology 54(5), 1477–1482.

Jin, H., Huo, L., Yang, Y., Lv, Y., Wang, J., Maillard, J., Holliger, C., Löffler, F.E. and Yan, J. 2023. Sulfurospirillum diekertiae sp. nov., a tetrachloroethene-respiring bacterium isolated from contaminated soil. International Journal of Systematic and Evolutionary Microbiology 73(2), 005693.

Johnson, D.B. and Sánchez-Andrea, I. 2019. Dissimilatory reduction of sulfate and zero-valent sulfur at low pH and its significance for bioremediation and metal recovery. Advances in Microbial Physiology 75, 205–231.

Kamyshny, A. 2009. Solubility of cyclooctasulfur in pure water and sea water at different temperatures. Geochimica et Cosmochimica Acta 73(20), 6022–6028.

Kamyshny, A., Gun, J., Rizkov, D., Voitsekovski, T. and Lev, O. 2007. Equilibrium distribution of polysulfide ions in aqueous solutions at different temperatures by rapid single phase derivatization. Environmental Science and Technology 41(7), 2395–2400.

Kassambara, A. 2020 ggpubr:“ggplot2” based publication ready plots.

Kumar, M., Nandi, M. and Pakshirajan, K. 2021. Recent advances in heavy metal recovery from wastewater by biogenic sulfide precipitation. Journal of Environmental Management 278, 111555–111555.

Lahti, L. and Shetty, S. 2017 Tools for microbiome analysis in R.

Lewis, A.E. 2010. Review of metal sulphide precipitation. Hydrometallurgy 104(2), 222–234.

McMurdie, P.J. and Holmes, S. 2013. Phyloseq: An R Package for Reproducible Interactive Analysis and Graphics of Microbiome Census Data. PLoS ONE 8(4), 1–11.

Mino, S., Kudo, H., Arai, T., Sawabe, T., Takai, K. and Nakagawa, S. 2014. Sulfurovum aggregans sp. nov., a hydrogen-oxidizing, thiosulfate-reducing chemolithoautotroph within the ε-proteobacteria isolated from a deep-sea hydrothermal vent chimney, and an emended description of the genus Sulfurovum. International Journal of Systematic and Evolutionary Microbiology 64(Pt_9), 3195–3201.

Miroshnichenko, M.L., Gongadze, G.A., Lysenko, A.M. and Bonch-Osmolovskaya, E.A. 1994. Desulfurella multipotens sp. nov., a new sulfur-respiring thermophilic eubacterium from Raoul Island (Kermadec archipelago, New Zealand). Archives of Microbiology 161(1), 88­93.

Miroshnichenko, M.L., Rainey, F.A., Hippe, H., Chernyh, N.A., Kostrikina, N.A. and Bonch-Osmolovskaya, E.A. 1998. Desulfurella kamchatkensis sp. nov. and Desulfurella propionica sp. nov., new sulfur-respiring thermophilic bacteria from Kamchatka thermal environments. International Journal of Systematic Bacteriology 48(2), 475–479.

Mori, K., Kim, H., Kakegawa, T. and Hanada, S. 2003. A novel lineage of sulfate-reducing microorganisms: Thermodesulfobiaceae fam. nov., Thermodesulfobium narugense, gen. nov., sp. nov., a new thermophilic isolate from a hot spring. Extremophiles 7(4), 283–290.

Ntagia, E., Chatzigiannidou, I., Williamson, A.J., Arends, J.B.A. and Rabaey, K. 2020. Homoacetogenesis and microbial community composition are shaped by pH and total sulfide concentration. Microbial biotechnology 13(4), 1026–1038.

Panda, S.K., Jyoti, V., Bhadra, B., Nayak, K.C., Shivaji, S., Rainey, F.A. and Das, S.K. 2009. Thiomonas bhubaneswarensis sp. nov., an obligately mixotrophic, moderately thermophilic, thiosulfate-oxidizing bacterium. International Journal of Systematic and Evolutionary Microbiology 59(9), 2171–2175.

Poncheewin, W., Hermes, G.D.A., van Dam, J.C.J., Koehorst, J.J., Smidt, H. and Schaap, P.J. 2020. NG-Tax 2.0: A semantic framework for high-throughput amplicon analysis. Frontiers in Genetics 10(January), 1–12.

Qiu, Y.Y., Guo, J.H., Zhang, L., Chen, G.H. and Jiang, F. 2017. A high-rate sulfidogenic process based on elemental sulfur reduction: Cost-effectiveness evaluation and microbial community analysis. Biochemical Engineering Journal 128, 26–32.

Quast, C., Pruesse, E., Yilmaz, P., Gerken, J., Schweer, T., Yarza, P., Peplies, J. and Glöckner, F.O. 2013. The SILVA ribosomal RNA gene database project: Improved data processing and web-based tools. Nucleic Acids Research 41(D1), 590–596.

Rittmann, B.E. and McCarty, P.L. (2020) Environmental Biotechnology: Principles and Applications, McGraw-Hill Education, New York.

Ross, D.E., Marshall, C.W., May, H.D. and Norman, R.S. 2016. Comparative genomic analysis of Sulfurospirillum cavolei MES reconstructed from the metagenome of an electrosynthetic microbiome. PLOS ONE 11(3), e0151214–e0151214.

Schauder, R. and Müller, E. 1993. Polysulfide as a possible substrate for sulfur-reducing bacteria. Archives of Microbiology 160(5), 377–382.

Schumacher, W., Kroneck, P.M.H. and Pfennig, N. 1992. Comparative systematic study on “Spirillum” 5175, Campylobacter and Wolinella species. Archives of Microbiology 158(4), 287–293.

Segerer, A., Stetter, K.O. and Klink, F. 1985. Two contrary modes of chemolithotrophy in the same archaebacterium. Nature 313(6005), 787–789.

Stetter, K.O. and Gaag, G. 1983. Reduction of molecular sulphur by methanogenic bacteria. Nature 305(5932), 309–311.

Steudel, R. (2020) Environmental technologies to treat sulfur pollution: Principles and engineering. Lens, P. and Pol, L.H. (eds), pp. 11-53, IWA Publishing, London, UK.

Stolz, J.F., Ellis, D.J., Blum, J.S., Ahmann, D., Lovley, D.R. and Oremland, R.S. 1999. Sulfurospirillum barnesii sp. nov. and Sulfurospirillum arsenophilum sp. nov., new members of the Sulfurospirillum clade of the ε-Proteobacteria. International Journal of Systematic and Evolutionary Microbiology 49(3), 1177–1180.

Sun, R., Li, Y., Lin, N., Ou, C., Wang, X., Zhang, L. and Jiang, F. 2020a. Removal of heavy metals using a novel sulfidogenic AMD treatment system with sulfur reduction: Configuration, performance, critical parameters and economic analysis. Environment International 136.

Sun, R., Zhang, L., Wang, X., Ou, C., Lin, N., Xu, S., Qiu, Y.Y. and Jiang, F. 2020b. Elemental sulfur-driven sulfidogenic process under highly acidic conditions for sulfate-rich acid mine drainage treatment: Performance and microbial community analysis. Water Research 185, 116230–116230.

Sun, R., Zhang, L., Zhang, Z., Chen, G.H. and Jiang, F. 2018. Realizing high-rate sulfur reduction under sulfate-rich conditions in a biological sulfide production system to treat metal-laden wastewater deficient in organic matter. Water Research 131, 239–245.

Takahashi, Y., Suto, K. and Inoue, C. 2010. Polysulfide reduction by Clostridium relatives isolated from sulfate-reducing enrichment cultures. Journal of Bioscience and Bioengineering 109(4), 372–380.

Takai, K., Kobayashi, H., Nealson, K.H. and Horikoshi, K. 2003. Deferribacter desulfuricans sp. nov., a novel sulfur-, nitrate-and arsenate-reducing thermophile isolated from a deep-sea hydrothermal vent. international journal of systematic and evolutionary microbiology 53(3), 839–846.

Thabet, O.B.D., Wafa, T., Eltaief, K., Cayol, J.L., Hamdi, M., Fauque, G. and Fardeau, M.L. 2011. Desulfovibrio legallis sp. nov.: A moderately halophilic, sulfate-reducing bacterium isolated from a wastewater digestor in Tunisia. Current Microbiology 62(2), 486–491.

Thompson, L.R., Sanders, J.G., McDonald, D., Amir, A., Ladau, J., Locey, K.J., Prill, R.J., Tripathi, A., Gibbons, S.M. and Ackermann, G. 2017. A communal catalogue reveals Earth’s multiscale microbial diversity. Nature 551(7681), 457–463.

Vésteinsdóttir, H., Reynisdóttir, D.B. and Örlygsson, J. 2011. Thiomonas islandica sp. nov., a moderately thermophilic, hydrogen- and sulfur-oxidizing betaproteobacterium isolated from a hot spring. International Journal of Systematic and Evolutionary Microbiology 61(1), 132–137.

Wickham, H. (2008) Elegant Graphics for Data Analysis: ggplot2.

Wickham, H., Averick, M., Bryan, J., Chang, W., McGowan, L., François, R., Grolemund, G., Hayes, A., Henry, L., Hester, J., Kuhn, M., Pedersen, T., Miller, E., Bache, S., Müller, K., Ooms, J., Robinson, D., Seidel, D., Spinu, V., Takahashi, K., Vaughan, D., Wilke, C., Woo, K. and Yutani, H. 2019. Welcome to the Tidyverse. Journal of Open Source Software 4(43), 1686–1686.

Wolfe, R.S. and Pfennig, N. 1977. Reduction of sulfur by Spirillum 5175 and syntrophism with Chlorobium. Applied and Environmental Microbiology 33(2), 427–433.

Yilmaz, P., Parfrey, L.W., Yarza, P., Gerken, J., Pruesse, E., Quast, C., Schweer, T., Peplies, J., Ludwig, W. and Glöckner, F.O. 2014. The SILVA and “all-species Living Tree Project (LTP)” taxonomic frameworks. Nucleic Acids Research 42(D1), 643–648.

Zhang, L., Lin, X., Zhang, Z., Chen, G.-H. and Jiang, F. 2018a. Elemental sulfur as an electron acceptor for organic matter removal in a new high-rate anaerobic biological wastewater treatment process. Chemical Engineering Journal 331, 16–22.

Zhang, L., Zhang, Z., Sun, R., Liang, S., Chen, G.H., Jiang, F., Lin, X., Zhang, Z., Chen, G.H. and Jiang, F. 2018b. Self-accelerating sulfur reduction via polysulfide to realize a high-rate sulfidogenic reactor for wastewater treatment. Water Research 130, 161–167.

Zhou, X., Liu, L., Chen, C., Ren, N., Wang, A. and Lee, D.J. 2011. Reduction of produced elementary sulfur in denitrifying sulfide removal process. Applied Microbiology and Biotechnology 90(3), 1129–1136.

