## Supplementary material 1 to 5, described in the text. for "Biological S^0^ reduction at neutral (pH 6.9) and acidic (pH 3.8) conditions: Performance and microbial community shifts in a H_2_/CO_2_-fed bioreactor"

### S.I.1. Reactor event log

**Table S.I. 1** Log summary of the events occurred during the operation of the gas-lift reactor and corrective actions taken.

| Operation day | Event | Corrective action |
| --- | --- | --- |
| 1 | Online pH drifted in 0.5 units of that of the offline measurements. | Probe was recalibrated, but the difference between online and offline measurements persisted. The pH control limits were modified to match the offline measurements. |
| 8 | Cloggage in the influent line due to granules accumulation in the influent entrance. | The lines were backflushed and contents returned to the reactor. Reactor was sealed and all influent and recirculation lines were replaced. Replaced lines were filled with anaerobic water to reduce/avoid oxygen introduction to the reactor. |
| 9 | Power failure during four hours. | All gas and liquid influent and recirculation systems were off during this period. The system was operated in batch during 3 hours after power restore. |
| 10 | Flow calibration of the influent pump lost. HRT increased to 3.6 days. | All liquid pumps were re-calibrated. |
| 13 | Online pH measurements started to drift to that of the offline measurements. Actual reactor pH 4.8. | The pH electrode was replaced with a new electrode. New electrode was calibrated and verified. |
| 23 | Power cut during 12 hours. | Planned maintenance of power lines. Heat exchanger units, gas recirculation, and influent media pumps turned off. Influent gas mass flow and pH controllers remained on with an alternate power supply. The reactor was operated in batch mode during this time. |
| 35 | Cloggage in the liquid recirculation line. Sulfur accumulation in the line. | Line flow backflushed. To contain backflush liquid leaving the reactor, the effluent line was closed. After the successful liberation of the line, the influent was turned off for 30 minutes to allow the solids to settle in the external settler. |
| 57 | Cloggage and rupture of the liquid recirculation line. Sulfur accumulation in the line. ~200 mL recirculation fluid lost. | Liquid recirculation line replaced. The line was filled with liquid from the settler to prevent oxygen from entering the reactor through the line. Recirculation rate was increased to 3.6 mL.min^-1^ to increase the solid flow in the line and avoid solid settlement in the line. |
| 59 | Decrease efficiency of suspended solids recovery. | Liquid recirculation rate decreased to 2.0 mL.min^-1^ |
| 64 | Gas effluent line partially clogged. | Carbonate accumulation in the effluent gas line. Line replaced and new sparger installed. |
| 66 | Cloggage and rupture of the liquid recirculation line. Sulfur accumulation in the line. No fluid lost. | Liquid recirculation line was replaced. The line was filled with liquid from the settler to prevent oxygen from entering the reactor through the line. |
| 69 | Decrease efficiency of suspended solids recovery. | As part of the corrective procedure on day 66, the liquid recirculation pump was left at 8.0 mL.min^-1^ by error. The flow was corrected to 2.0 mL.min^-1^. |
| 83 | Gas effluent line partially clogged. Gas partially leaving the through liquid effluent. | Carbonate accumulation in the effluent gas line. Line replaced and new sparger installed. |
| 113 | Damage of the pump head of the liquid recirculation line. | No flow of liquid through the pump. The pump head was replaced. The line was filled with liquid from the settler to prevent oxygen from entering the reactor through the line. |

### S.I.2. Thiosulfate formation

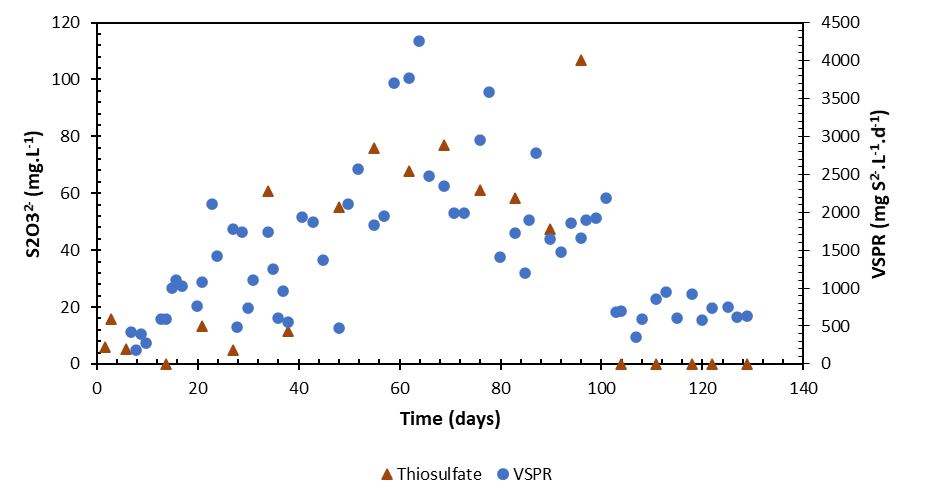

**Figure S.I. 1** Concentration of dissolved thiosulfate (red triangles) contrasted with the volumetric sulfide producing rates (VSPR) in the 4 L gas lift rector.

### S.I.3. Volatile fatty acid (VFA) composition

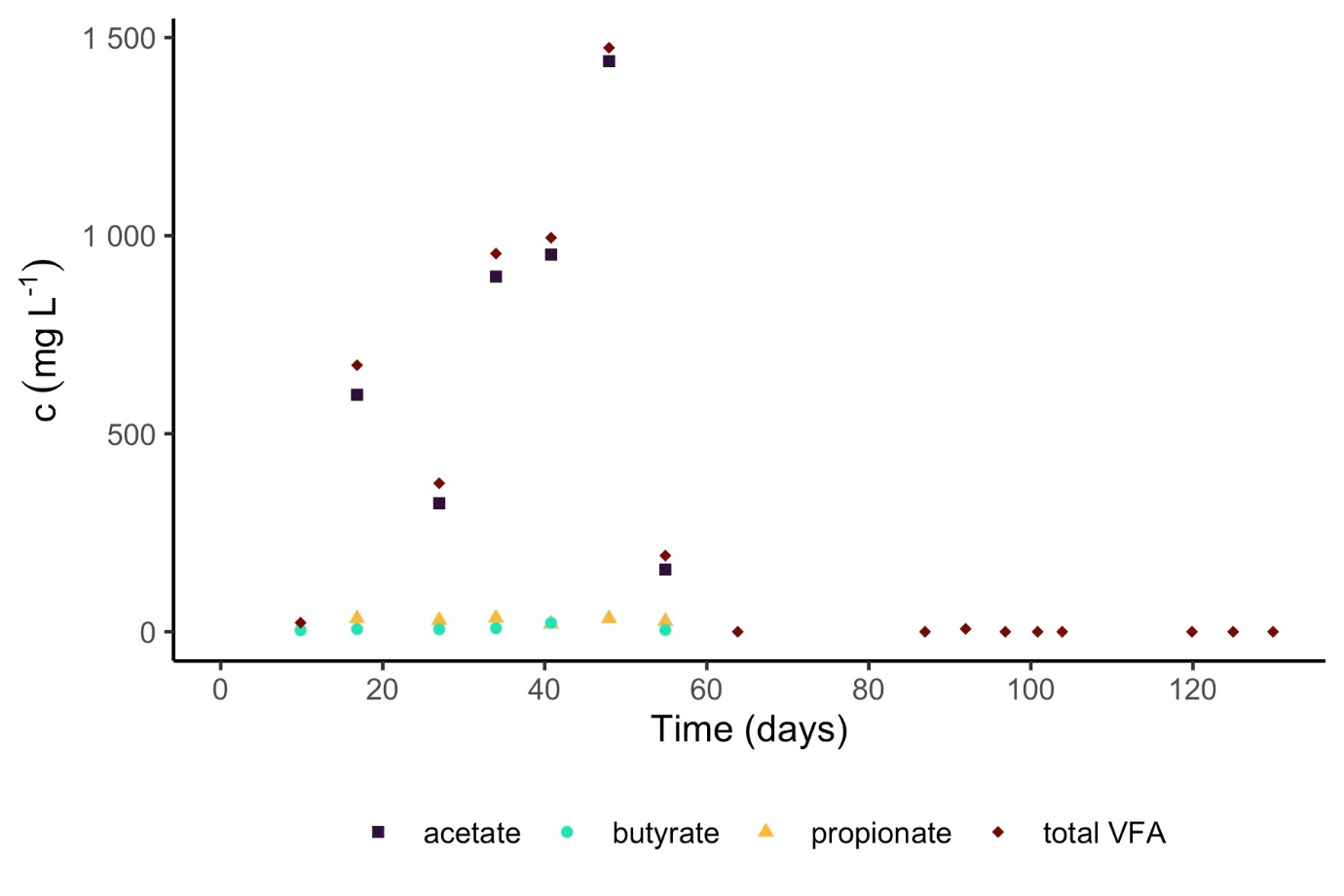

**Figure S.I. 2** Fraction of the volatile fatty acid (VFA) found in the gas-lift samples. In the plot are shown only the VFA fraction found in the samples. VFA fractions measured under the limit of quantification are not shown.

### S.I.4. 16S rRNA gene amplicon sequence reads

**Table S.I. 2** sequencing depth per sample after quality control and filtering

| **Sample** | **Reads** |
| --- | --- |
| biosulfur_A | 599508 |
| biosulfur_B | 100273 |
| biosulfur_C | 233324 |
| Emm_20200306_A | 561669 |
| Emm_20200306_B | 480109 |
| Emm_20200306_C | 143427 |
| Emm_20210604_A | 637884 |
| Emm_20210604_B | 360171 |
| Emm_20210604_C | 22058 |
| Negcontrol_A | 581 |
| R2_d24_A | 350499 |
| R2_d24_B | 471115 |
| R2_d24_C | 449994 |
| R2_d59_A | 305673 |
| R2_d59_B | 614855 |
| R2_d59_C | 529843 |
| R2_d73_A | 398199 |
| R2_d73_B | 522318 |
| R2_d73_C | 524306 |
| R2_d101_A | 696963 |
| R2_d101_B | 462725 |
| R2_d101_C | 421098 |
| R2_d118_A | 595869 |
| R2_d118_B | 560330 |
| R2_d118_C | 393985 |
| R2_d130_A | 152177 |
| R2_d130_B | 84542 |
| R2_d130_C | 254896 |

### S.I.5. Relative read abundance of top 50 most abundant taxa

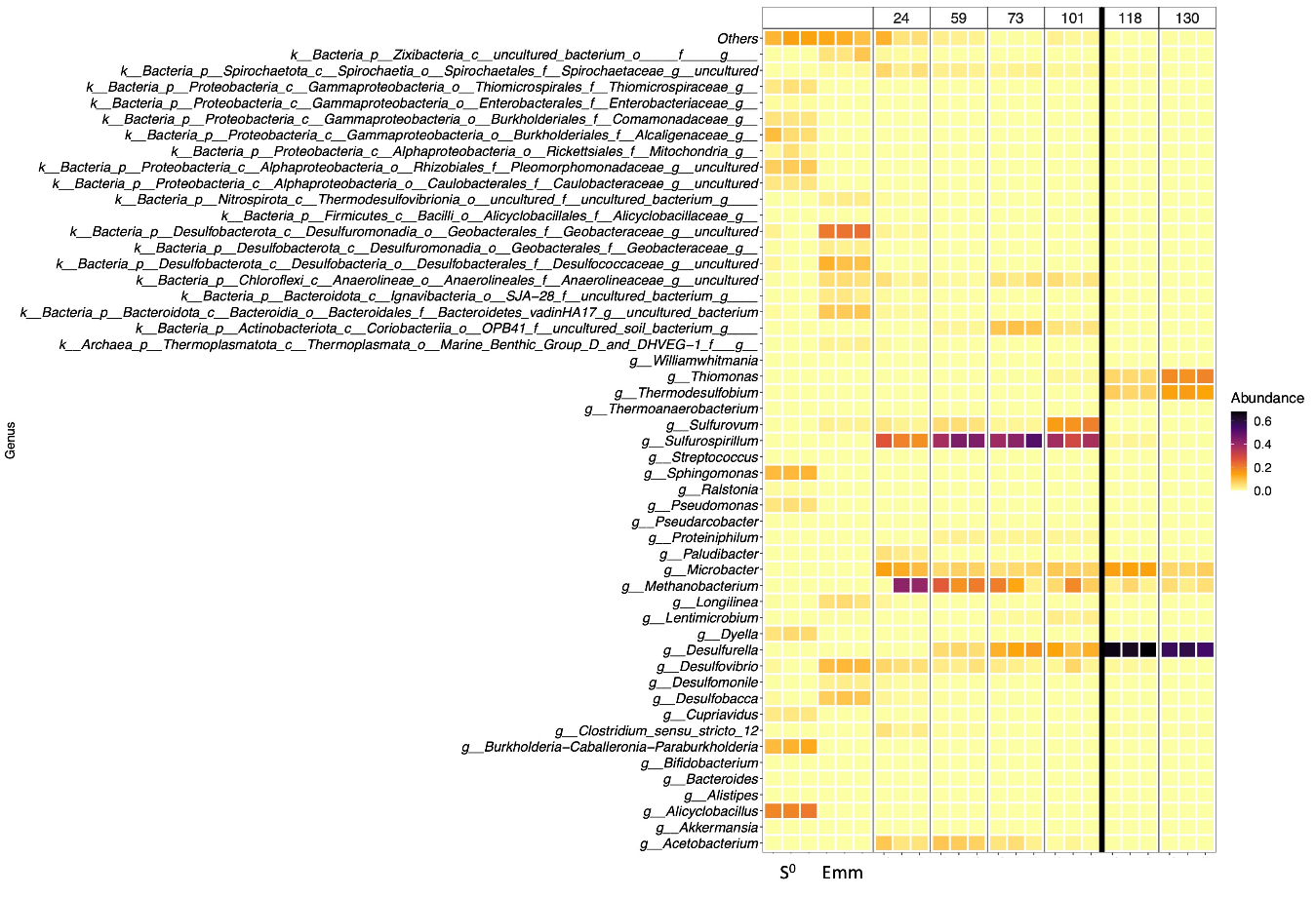

**Figure S.I. 3** Top 50 most abundant taxa detected in samples from the original anaerobic sludge used as inoculum (Emm), the S^0^, and the reactor operated at pH 6.9 (days 24 – 101) and pH 3.9 (day 118 and 130), separated by the black line.

### S.I.6. Microscopy images

**A.**

| 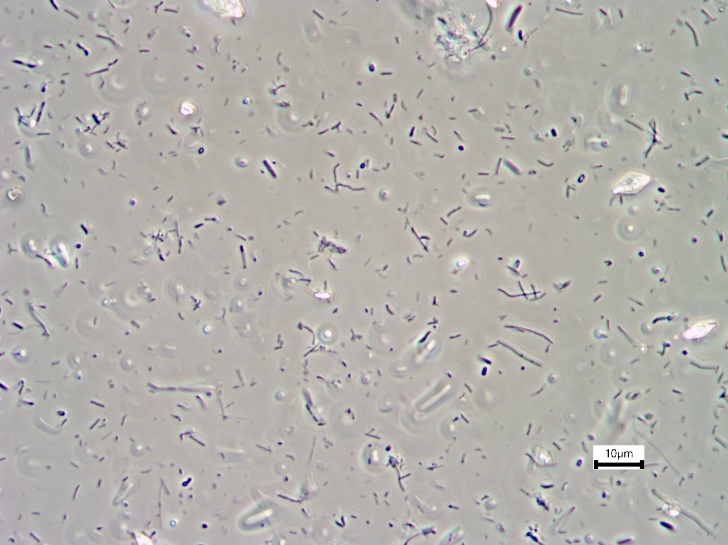 | 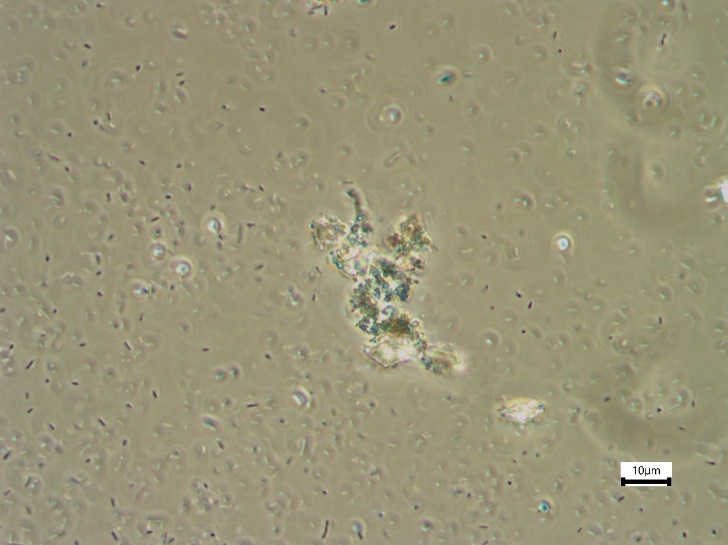  **B.**  **C.** |
| --- | --- |
| 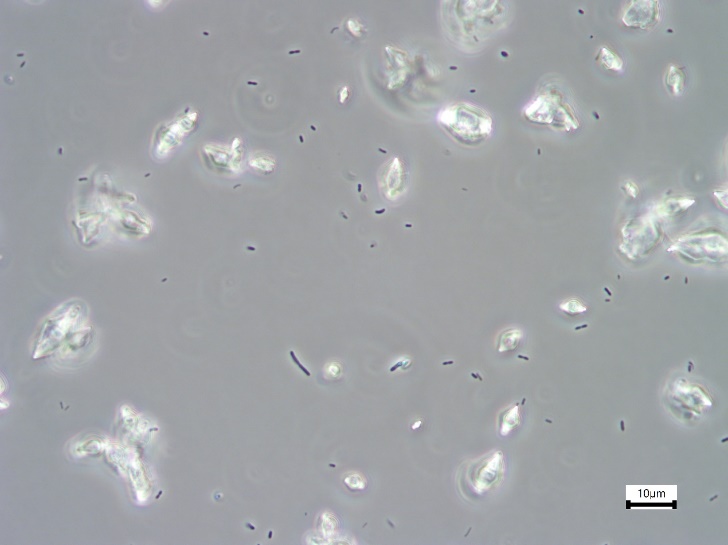 | |

***Figure S.I. 4*** *Contrast microscopy images of the gas lift reactor samples on operational days A. 20, B. 92 (steady-state neutral regime), and C. 120 (steady-state acidic regime).*
